# Improving design and normalization of multiplex proteomics study

**DOI:** 10.1101/2024.12.05.627093

**Authors:** Huaying Fang, Mei-Chiung Shih, Lihua Jiang, Felipe da Veiga Leprevost, Ruiqi Jian, Joanne Chan, Alexey I. Nesvizhskii, Michael P. Snyder, Hua Tang

## Abstract

Advances in multiplex mass spectrometry-based technologies have enabled high-throughput, quantitative proteome profiling of large cohort. However, certain experimental design configurations can amplify sample variability and introduce systematic biases. To address these challenges, we incorporated two novel features in a recent proteogenomic investigation: (1) the inclusion of two reference samples within each mass spectrometry run to serve as internal standards, and (2) the analysis of each specimen as technical replicates across two distinct mass spectrometry runs. Building on these enhancements, we present ProMix, a flexible analytical framework designed to fully leverage these supplementary experimental components. Using both simulated and real-world datasets, we demonstrate the improved performance of ProMix and highlight the advantages conferred by these refined experimental design strategies.

## Introduction

Multiplexed quantitative mass spectrometry (MS)-based proteomics approaches, such as Tandem Mass-Tag (TMT) and multiplexed isobaric tags for relative and absolute quantitation (iTRAQ), enable comprehensive and multifaceted interrogation of the proteomes in biofluid, tissues and cells^1,2^. Achieving reliable data quality at an expanding throughput, these approaches have been widely adopted in large-scale clinical and proteogenomic studies^3–5^.

The key component of TMT and iTRAQ proteomic approaches is the use of isobaric tags, which have identical masses but feature distinct isotopes. In a typical mas-spectrometry run (hereafter abbreviated as a “*run*”), peptides in each sample are labelled with a unique isobaric tag (Fig 1A). Tagged peptides from multiple samples are mixed and go through liquid chromatography (LC) followed by tandem spectrometry. Peptide and protein identification can proceed following data-dependent (DDA) or data-independent acquisition approach (DIA). Peptides labeled with different tags behave indistinguishably during LC and mass-analysis; however, the tag-specific isotopes are revealed after peptide fragmentation, and the intensity ratio of the TMT-tag are used as quantitative readouts (Fig 1B). In studies that perform multiple runs, a *reference* sample is included in every run and serves as an internal standard, facilitating data integration and comparison across all runs.

**Figure 1.**
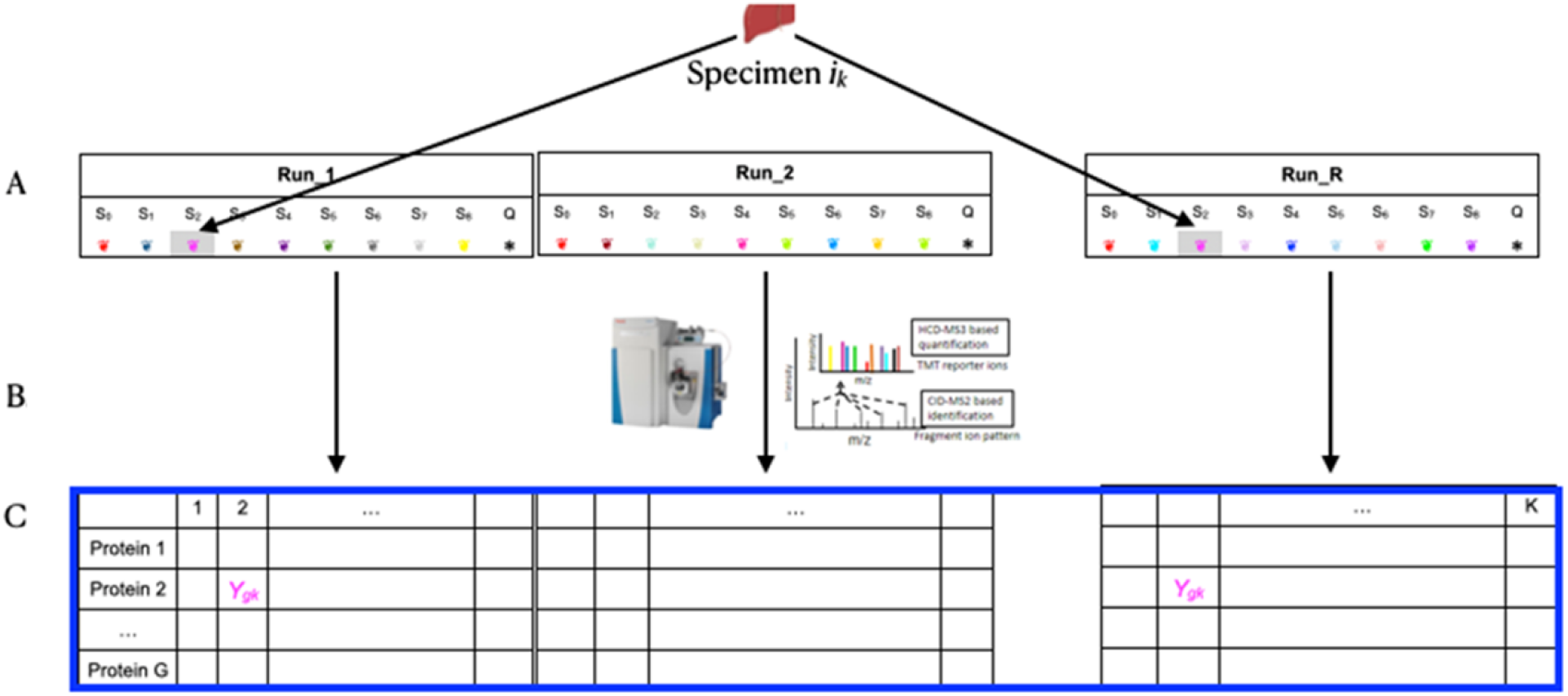
Experimental design and data representation of the proteomics study of GTEx lung tissues. (A) Peptides extracted from tissue donors were grouped into mass-spectrometry runs and labeled by TMT-tags. Each run included a common reference (denoted by Q), a mega-pool (denoted by S0) and eight lung samples (denoted by S1, …, S8). Each specimen was analyzed in two distinct runs; samples S2 in Run_1 and Run_R illustrate a pair of technical replicates derived from the same tissue specimen. (B) TMT-LC-MS platform generates spectra that allow for both the identification of the protein and the quantification of the relative abundance of the protein in each of the samples relative to the reference. (C) The relative abundance of a protein in a sample is defined as the ratio of the protein in the sample to that in the tissue-specific reference from the same run and is denoted as 𝑌_𝑔𝑘_.

In such cases, protein quantification, represented as the log ratio of relative abundance between a sample and the reference, requires normalized prior to downstream analyses. Probability quotient normalization (PQN), defined in the next section, is a simple, effective and widely adopted normalization procedure^6^. PQN adjusts the log ratios of all proteins in a sample by subtracting a constant value such that the median log ratio across all proteins is zero in each sample. Thus, this method accounts for stochastic variations in the absolute total protein quantities in input samples, assuming the protein composition is similar across all samples.

However, PQN does not account for variability in the measurements of the reference samples across runs. Such variation in the reference samples not only adds noise to each sample, but also induces an undesired dependent structure within a run, which is difficult to model and complicates downstream analysis.

Our work is motivated by ongoing proteogenomic studies, and we propose a normalization procedure that leverages two key design features, a second common reference in each run and replicates of each biological specimen. The proposed statistical model, ProMix, jointly accounts for the variability in the total absolute protein amount of each sample and the uncertainty associated with each protein measurement in the reference sample. The performance of ProMix, as well as the benefits of including a second reference and replicate samples, are evaluated using simulated and real data. These analyses demonstrate that the combination of ProMix and experimental design considerations can substantially improve the precision and consistency of protein quantification across runs. ProMix is implemented in R and is available at https://github.com/tanglab/ProxMix.

## Methods

### Motivating example: experimental setup, data and notations

We begin by describing the study that will serve as a data example in this paper, which aims to quantify protein abundance in 144 lung tissues from the Gene-Tissue-Expression consortium (GTEx) using a 10-plex TMT-LC-MS platform. Details about the GTEx project and sample collection can be found in previous publications^7,8^. In each run, eight of the ten TMT-labelled samples represent peptides derived from individual donors. The remaining two tags represented two distinct pooled samples used as internal standards: a tissue-specific reference consisted of the peptide mixture of all lung specimens analyzed, and a mega-pool, composed of peptides derived from 201 tissue specimens representing 32 tissue sites from a previous study^8^. In each run, approximately 15*ug* of multiplexed peptide mixture was fractionated using two-dimensional reversed-phase liquid-chromatography. The first dimension used high-pH separation, followed by orthogonal low-pH separation in the second dimension. Mass-spectrometry data were acquired using two Thermo-Fisher Orbitrap Fusion Lumos instruments. In total, 36 runs were performed. Peptide and protein identification and quantification were performed using MSFragger and Philosopher as previously described ^9–11^. Peptides from each specimen were labeled and split into two equal aliquots. These technical replicates were analyzed in two different runs, one on each instrument. An example of a technical replicate pair is illustrated in Fig 1A. For clarity, we use the term “specimen” to refer to biological tissues, while reserving the term "sample" to indicate single replicates.

Following common practice, we use the relative abundance as the basic input data, computed as the log2 ratio of a protein in a sample relative to that in the tissue-specific reference measured in the same run. Henceforth, this quantity will be abbreviated as “abundance.” The abundance data from a run are represented as a matrix, with each row representing a protein, and each column representing a sample. Subsequently, data from all runs are concatenated to form the input data matrix, 𝑌_𝑔𝑘_, in which *g=1, …, G* indexes the union of proteins observed across all runs, and *k=1, …, K* denotes the sample index (Fig 1). In the lung study, K=324, including 144×2=288 samples from lung donors and 36 mega-pools. In total G=9063 proteins were detected in at least one of the 288 lung samples. We note that because some proteins are quantified only in certain samples or runs, the matrix 𝑌_𝑔𝑘_ typically contains a substantial amount of missing data.

For convenience, we will use the notation 𝑙_𝑘_ ∈ {0,1, … , 𝐿 = 144} to map the sample index *k* to the specimen index, 𝑙, with 𝑙_𝑘_ = 0 for mega-pool and 1 ≤ 𝑙_𝑘_ ≤ 𝐿 for the 144 lung specimens. Likewise, we define 𝑟_𝑘_ ∈ {1,2, … , 𝑅 = 36} and 𝑡_𝑘_ ∈ {0,1, … , 8} to facilitate the mapping from the sample index to each of the 36 runs and to each of the 9 tag positions within a run, respectively. In each run, 𝑡_𝑘_ = 0 is reserved for the mega-pool.

### ProMix

With these notations, we propose a linear mixed model for protein abundance,

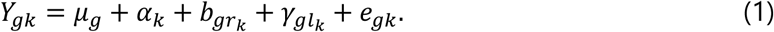

The definitions and interpretations of each term in the model are as follows. The term, 𝜇_𝑔_ denotes the overall deviation across all samples from the reference for protein 𝑔. Next, the term 𝛼_𝑘_ describes the deviation of sample *k* from the run-specific reference across all proteins. This term is a scale factor needed to account for the uncontrollable variation in the total absolute materials, also referred to as the dilution factor, in a sample. The term, 𝑏_𝑔𝑟𝑘_, represents the run-specific effect of protein 𝑔. Intuitively, this term can be interpreted as the measurement error of the protein in the reference in run, 𝑟_𝑘_ ∈ {1, … , 𝑅}. Recall that the abundances of a protein in all samples assayed in a run 𝑟_𝑘_ are computed relative to a common reference, and thus variability in the reference itself will induce a run-level correlation among the samples. For this reason, we model 𝑏_𝑔𝑟_ as a random effect: 𝑏_𝑔𝑟_ ∼N(0, 𝜙_𝑔_). The term 𝛾_𝑔𝑙_ is the variable of primary interest and represents the specimen effect of protein *g* in specimen 𝑙_𝑘_. The estimate 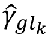 can be used as the normalized protein abundance in subsequent analyses. Finally, 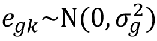 is the residual term for gene 𝑔 in sample 𝑘. Taken together, the parameters in the models are Ω =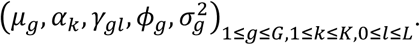

Directly estimating all parameters simultaneously is computationally expensive. Therefore, we implemented a coordinate-wise descent method, iterating through two steps. The computational algorithm of the proposed method, **ProMix**, is described in **Box 1**. Note that in Step (2.1), the update of (𝛼_𝑘_) can be computed for each sample, *k*, separately. Likewise, in Step

(2.2), 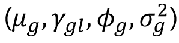 and (𝑏_𝑔𝑟_) can be updated for each protein, *g*, separately. This greatly

reduces the dimensionality of the parameter space and simplifies the optimization.

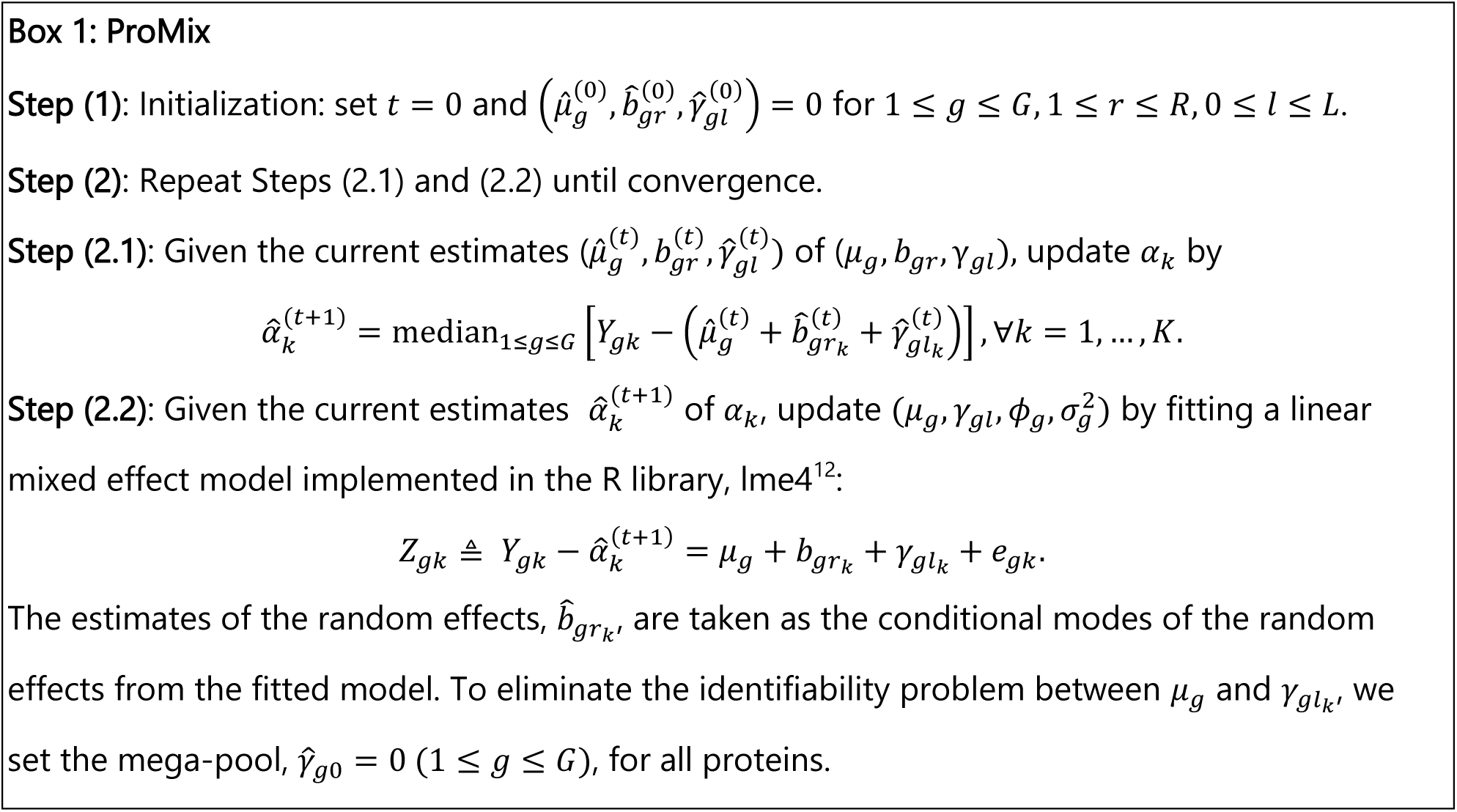

### PQN

The probability quotient normalization (PQN) is widely used and will serve as the primary alternative method for assessing the performance of ProMix. In PQN, the median log ratio between a sample relative to the reference is set to 0 for each sample:

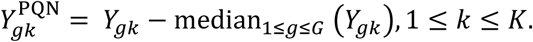

Note that 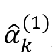, computed by Step (2.1) in the first iteration of ProMix is, in fact, the PQN normalization constant. Typically, the median is computed over all non-missing proteins in sample *k*; however, the missingness of a protein in a sample itself depends not only on its true abundance, but also the total abundance of the protein across all samples in a run, as well as the stochasticity of detection in a specific run. Therefore, to reduce the influence of these variably missing proteins, we compute the median over a subset of common proteins in simulation and real data analysis described below, defined as proteins with reference intensity greater than 20 and with no missing values in any of the mega-pool samples.

### Simulation of TMT data

The simulation is designed to evaluate normalization methods and experimental design features, while maintaining realistic complexities of the real dataset. Therefore, we simulated a dataset with 8000 proteins according to the additive model in Equation (1) and featuring a range of absolute, as well as relative, variation. The magnitude and variance for the model parameters were taken from the estimates in the real data from the lung tissue study. In addition, for 403 of these proteins, we set the run effects to 0 across all runs to evaluate the normalization methods in a setting, where PQN suffices and the random-effects term in ProMix, 𝑏_𝑔𝑟_, is unnecessary.

### Methods of assessment

For downstream analyses, the primary parameter of interest in equation (1) is 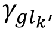 which can be interpreted as the normalized protein abundance of protein 𝑔 in specimen 𝑙_𝑘_. We use two metrics to assess the precision of the estimated protein abundance.

### RMSE

In simulated data, for which the true abundance is available, we can compare the square root of mean squared errors (RMSE) between the estimated and the true values,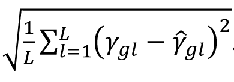 Since the mean (relative) abundance of each protein across all samples is arbitrary and plays no role in downstream analysis, we center the true and estimated mean abundance of each protein at 0 for all methods.

### Replication-RMSE (repRMSE)

As an alternative metric, we compare the RMSE between technical replicates, as these quantities should be reduced after adjustment of sample effects 𝛼_𝑘_ and run effects, 𝑏_𝑔𝑟𝑘_ . Because the repRMSE does not require the knowledge of the true values of 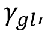 it can be used to evaluate normalization methods for both simulated and real data examples.

However, directly comparing repRMSE between ProMix and PQN gives ProMix an unfair advantage as ProMix uses the replicate information in estimating 𝑏_𝑔𝑟𝑘_ and 𝛾_𝑔𝑙𝑘_. In other words, the model can be overfitted and underestimates 𝑒_𝑔𝑘_. To overcome this caveat, we apply ProMix in a leave-one-out mode (ProMix-LOO), in which ProMix is applied to a subset of the data, leaving out one sample from each run. The run-specific parameters, 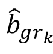, are then plugged in for the left-out samples. The repRMSE is only computed for replication pairs using the left-out samples. In both the real lung study and simulation, the two technical replicates of a specimen always have the same TMT-tag by design. This feature allows us to divide the full data into eight disjoint left-out subsets, each defined by samples with a specific TMT-tag 𝑡^∗^ ∈ {1, … 8}. We first demonstrate the validity of the leave-one-out procedure on the simulated data and then use the replication RMSE computed using ProMix-LOO to evaluate the normalization method on real data.

## Results

### Description of simulated data

We simulated a dataset with 8000 proteins. The median proportions of variance attributed to donor, mass-spec runs, sample, and random measurement errors are 0.43, 0.18, 0.07 and 0.21, respectively. These relative proportions are described in Fig 2.

**Figure 2.**
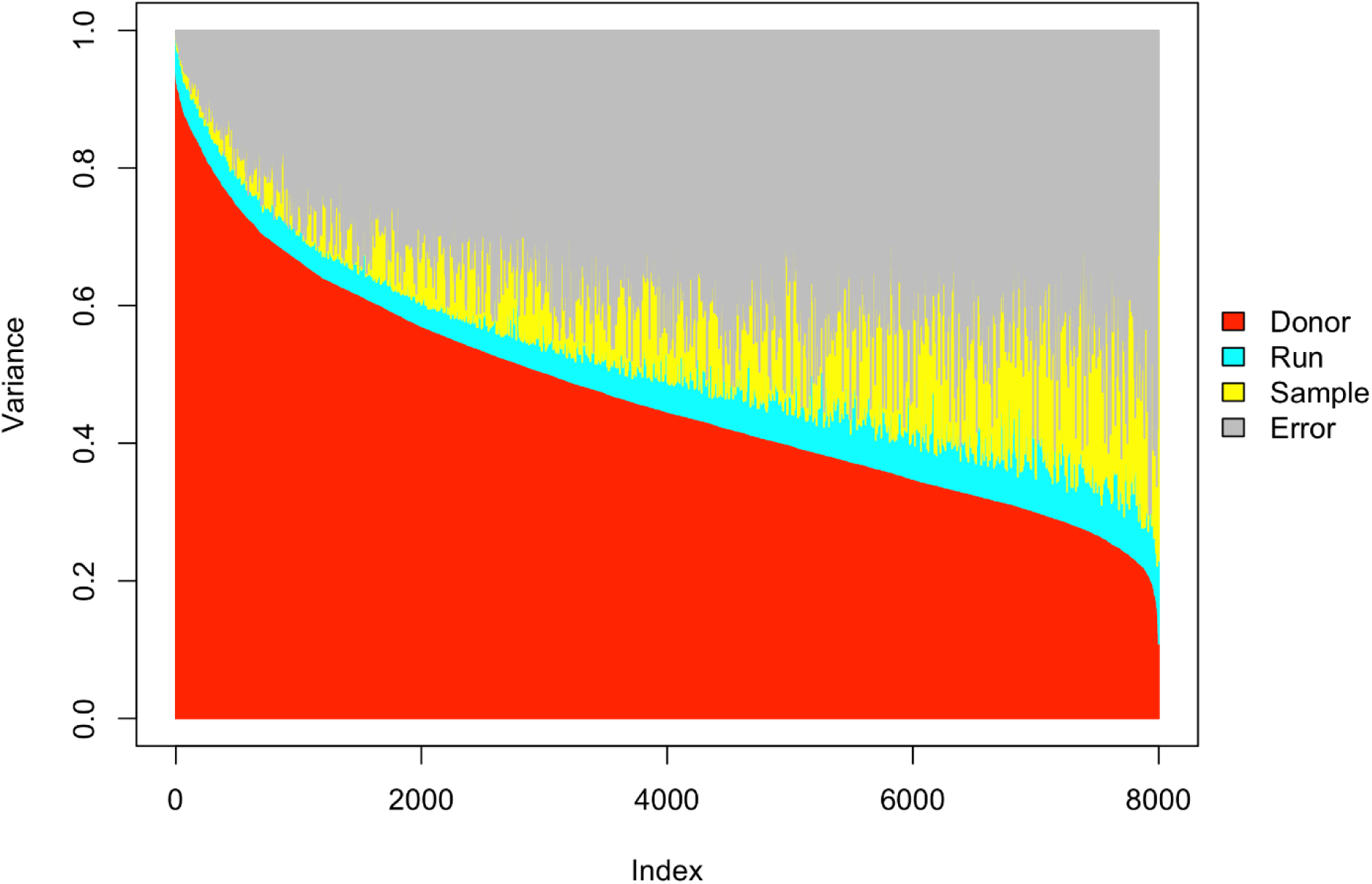
Description of simulated mass-spec protein quantification data. Each vertical segment represents a protein, in which the color of the segment corresponds to the relative proportions of variance attributed to donor (red, 𝛾_𝑔𝑙_ ), mass-spectrometry runs (cyan, 𝑏_𝑔𝑟_ ), sample (yellow, 𝛼_𝑘_), and random measurement errors (gray, 𝑒_𝑔𝑘_), respectively.

### Simulation Results

Fig 3 displays the accuracy of normalization, measured by the RMSE between the estimated and true donor-specific abundance. Both PQN and ProMix produced more accurate estimates of the donor effects compared to simply averaging the replicate measurements without normalization.

**Figure 3.**
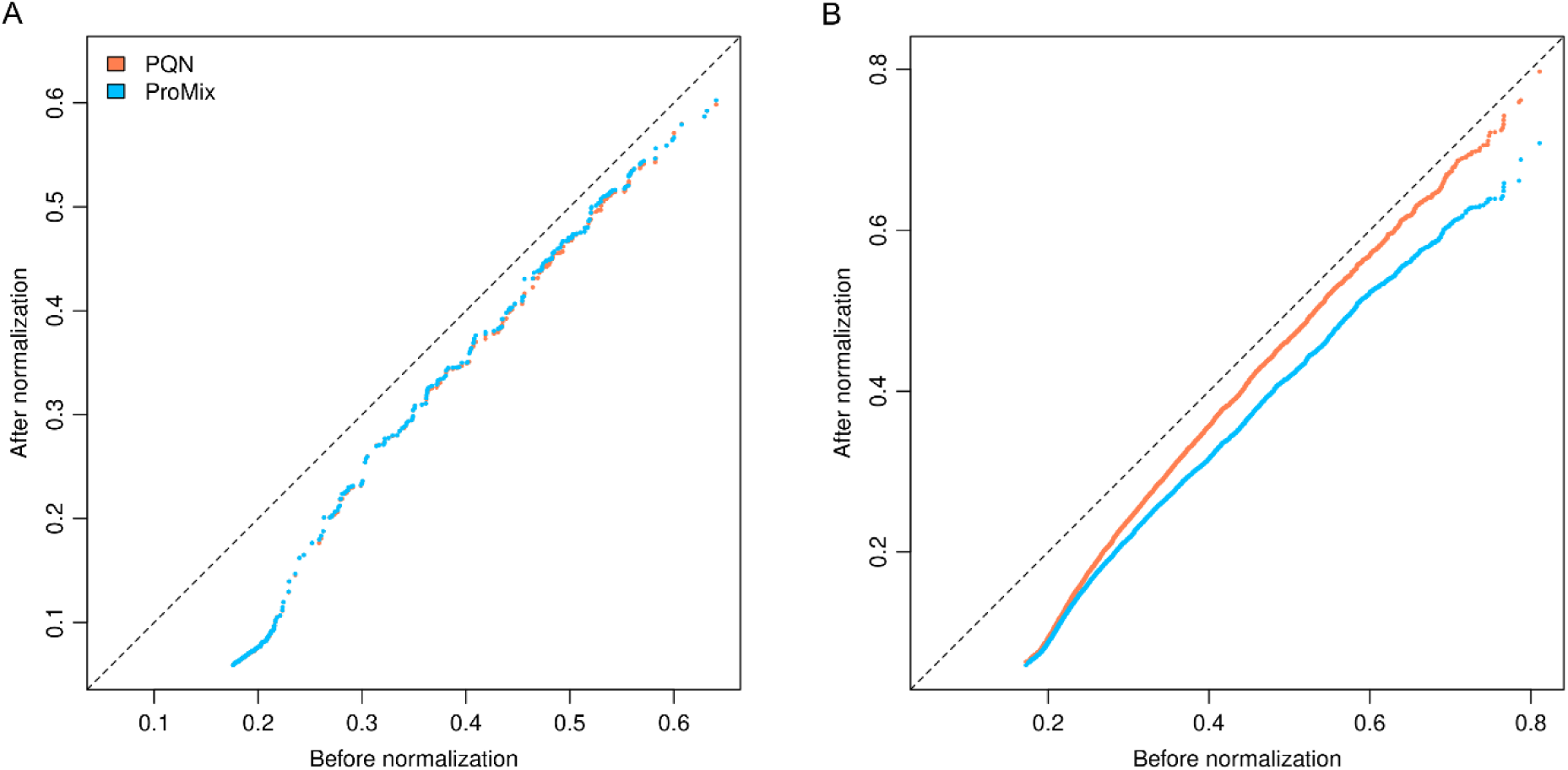
QQ-plot comparing the root mean squared error (RMSE) between the estimated and true donor effects on simulated data, before (x-axis) and after normalization (y-axis). (A) proteins whose true run-specific effect is 0 (e.g. 𝜙_𝑔_ = 0). (B) proteins whose true run-specific effect is not 0 (e.g. 𝜙_𝑔_ > 0).

Fig 3A describes simulated proteins with no run-specific effect: 𝜙_𝑔_ = 0. In this setting, PQN is the most parsimonious model and ProMix incurs a slight loss of efficiency by estimating the unnecessary run effects parameters. However, the loss of efficiency is minor, with PQN and ProMix achieving lower RMSE in 54% and 46% of the 403 proteins, respectively. In contrast, in the presence of run-specific effects, ProMix outperforms PQN for 93% of proteins, achieving a substantial reduction in RMSE (median 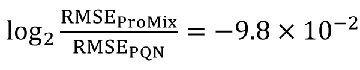; two-sided Wilcoxon test 𝑝 < 1 × 10^−56^, Fig 3B).

Likewise, using repRMSE as a measure of accuracy, which does not require the knowledge of the true protein abundance, reaches the same conclusion. For proteins without run effects, ProMix- LOO and PQN achieve similar accuracy as expected (Fig 4A). In contrast, when the run-specific effect is present, ProMix-LOO substantially outperforms PQN, achieving a lower repRMSE in 91% of the proteins (Fig 4B). The validity of using repRMSE as a surrogate measure for RMSE and the necessity of computing repRMSE in ProMix-LOO mode are demonstrated in Fig S2. Comparing ProMix-LOO with PQN, the reduction in repRMSE is highly predictive of the reduction in RMSE (𝑟^2^ = 0.65, 𝑝 < 2 × 10^−16^); in other words, the reduction in repRMSE using ProMix-LOO can serve as an informative surrogate for the reduction in RMSE. In contrast, the boxplots shown in the lower panel of Fig S2 illustrate that, using full data, ProMix appears to reduce repRMSE even for proteins without run-specific effects. For these proteins, ProMix reduces repRMSE without improving RMSE. Taken together, these results demonstrate that, for real data where the true protein abundance is unavailable, repRMSE offers an informative criterion for comparing across normalization methods and experimental design features, provided that ProMix is run in LOO mode.

**Figure 4.**
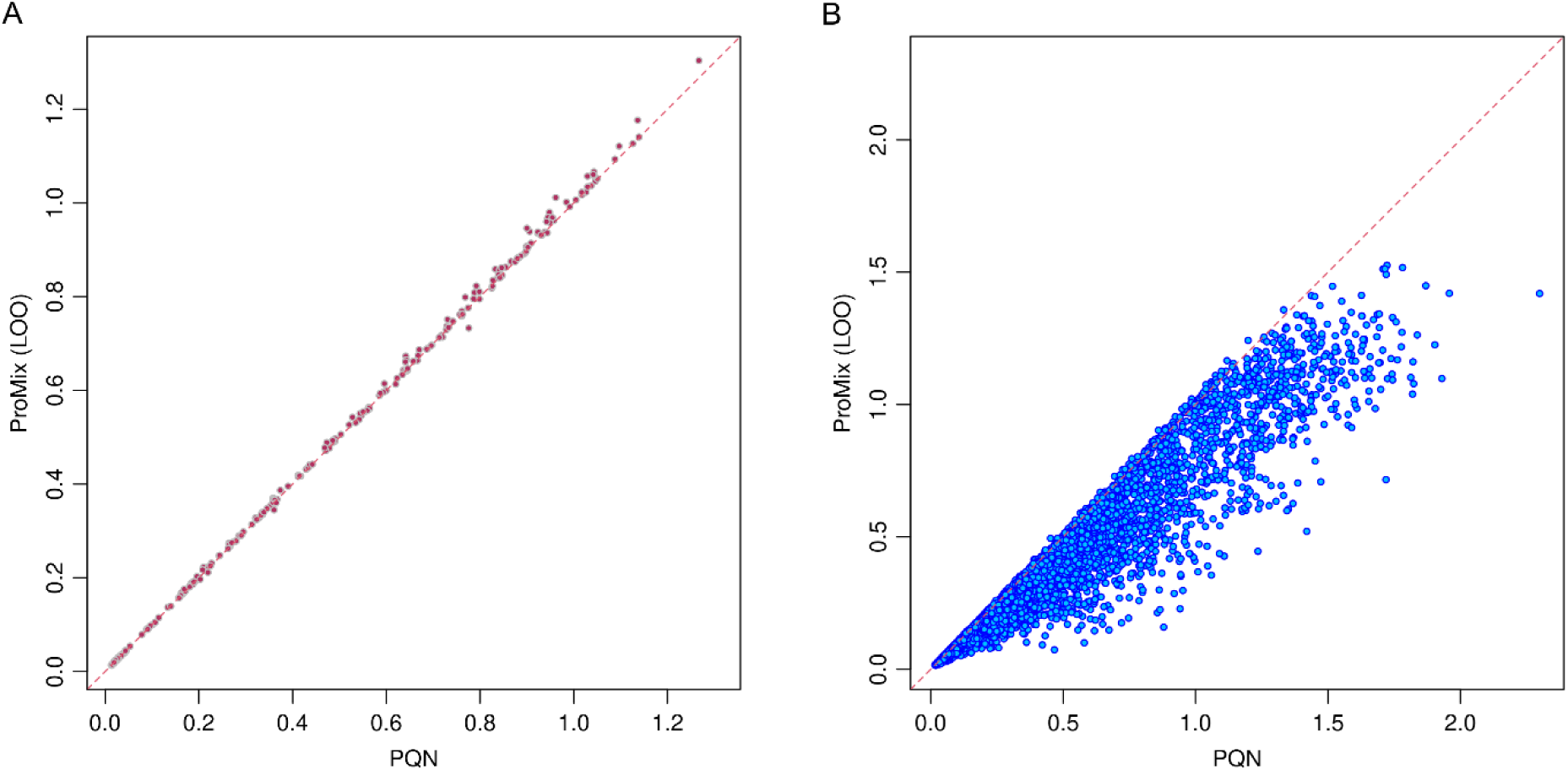
Comparisons of replication mean squared errors (repRMSE) in simulated data, using PQN (x-axis) or ProMix-LOO normalization (y-axis). (A) proteins whose true run-specific effect is 0 (e.g. 𝜙_𝑔_ = 0). (B) proteins whose true run-specific effect is not 0 (e.g. 𝜙_𝑔_ > 0).

### Design Considerations

We next use simulated data to examine the benefits of the inclusion of technical replicates and a second internal standard. This setup, used in the GTEx lung study, will be referred to as the “full design.” We consider three alternative experimental designs: (1) “SingleRef,” which includes technical replicates but not mega-pool; (2) “NoRep,” which includes two references in each run, but each specimen is analyzed in only one run; and (3) “SingleRef+NoRep,” which includes neither mega-pool nor replicates. This last design serves as a baseline and represents the standard practice in large-scale quantitative proteomic studies. While the model in Equation (1) is estimable under the SingleRef design, it is over-specified for the other two alternative designs. Therefore, for the NoRef design, we run ProMix with a reduced model, such that the run effects are estimated based on the mega-pool alone:

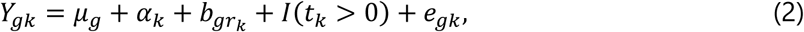

where 𝐼(⋅) is the indicator function. Finally, for “SingleRef+NoRep,” the run effects and specimen effects are unidentifiable without further assumption; hence we use PQN. Fig 5 compares the RMSE of the three alternative designs (y-axis) using the full design (x-axis) as the benchmark. For proteins without run effects, the SingleRef design achieves nearly identical performance to the full design; both designs outperform the other two alternatives that do not include replicates, NoRep and SingleRef+NoRep. This is not surprising: for these proteins, the estimation of run effects is not necessary, and the performance gain in both the full and SingleRef designs is primarily derived from the repeated measures of each specimen. In contrast, when run effects are present, including either technical replicates or a second internal standard in each run achieves substantial improvement over the baseline design. It is worth noting that the SingleRef design, which includes only technical replicates and not a second reference, performs nearly as well as the full design, suggesting that run effects can be estimated using technical replicates alone. On the other hand, when technical replicates are not available, including a second internal standard is an efficient alternative.

**Figure 5.**
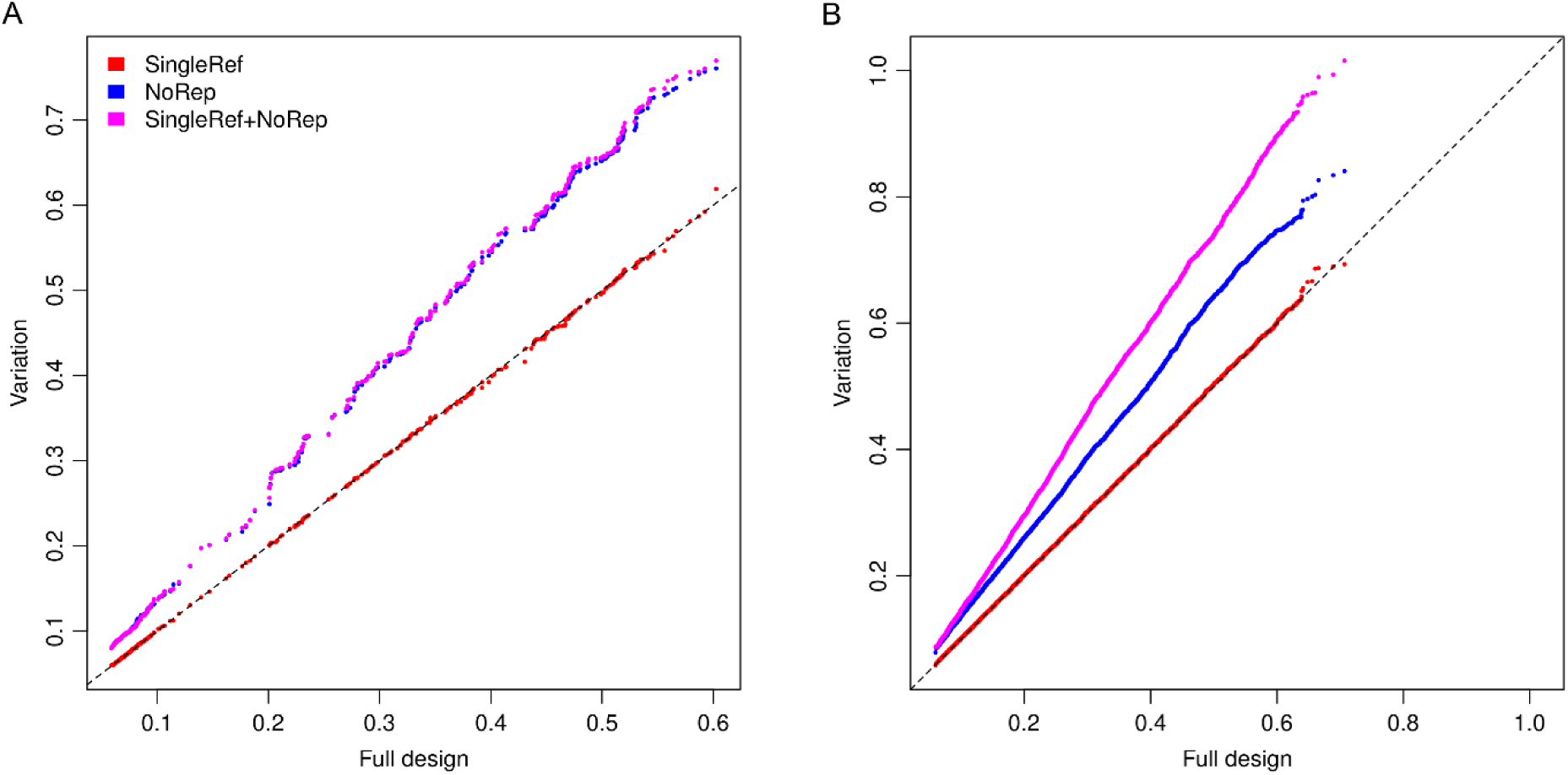
**Comparison of different experimental designs on simulated data**. (A) QQ-plot of proteins whose true run-specific effect is 0 (e.g. 𝜙_𝑔_ = 0). (B) QQ-plot of proteins whose true run-specific effect is not 0 (e.g. 𝜙_𝑔_ > 0). For both panels, the x-axis represents the MSE achieved in the full experimental design, in which each specimen is analyzed in two different runs, and each run includes two references. The y-axis represents the MSE achieved by variations of experimental designs: "SingleRef" includes technical replicates but only one reference in each run; "NoRep" includes two references in each run, but each specimen is analyzed in only one run; "SingleRef+NoRep" includes only one reference in each run, and each specimen is analyzed only in one run.

### Analysis of GTEx Lung Data

The lung dataset represents specimens from 144 distinct GTEx donors. Each specimen was analyzed in two runs to alleviate the missing data problem. In total, 9063 proteins were quantified in at least one specimen, with 7842 proteins measured in at least 72 (50%) of the specimens at least once. In contrast, 7088 proteins would be expected to reach the 50% non-missing rate without replicates. A median of 6937.5 proteins were quantified in each sample, and 4231 proteins were quantified in all 288 samples.

The distribution of missing proportions is depicted in Fig S1, which supports the observation that the missing pattern is correlated between samples within a run. We applied ProMix to 7762 proteins, which were quantified in at least 7 of the 36 runs and included at least 10 replicate pairs. Compared to PQN, ProMix-LOO achieves lower repMSE for 7233 (93%) proteins (Fig 6).

**Figure 6.**
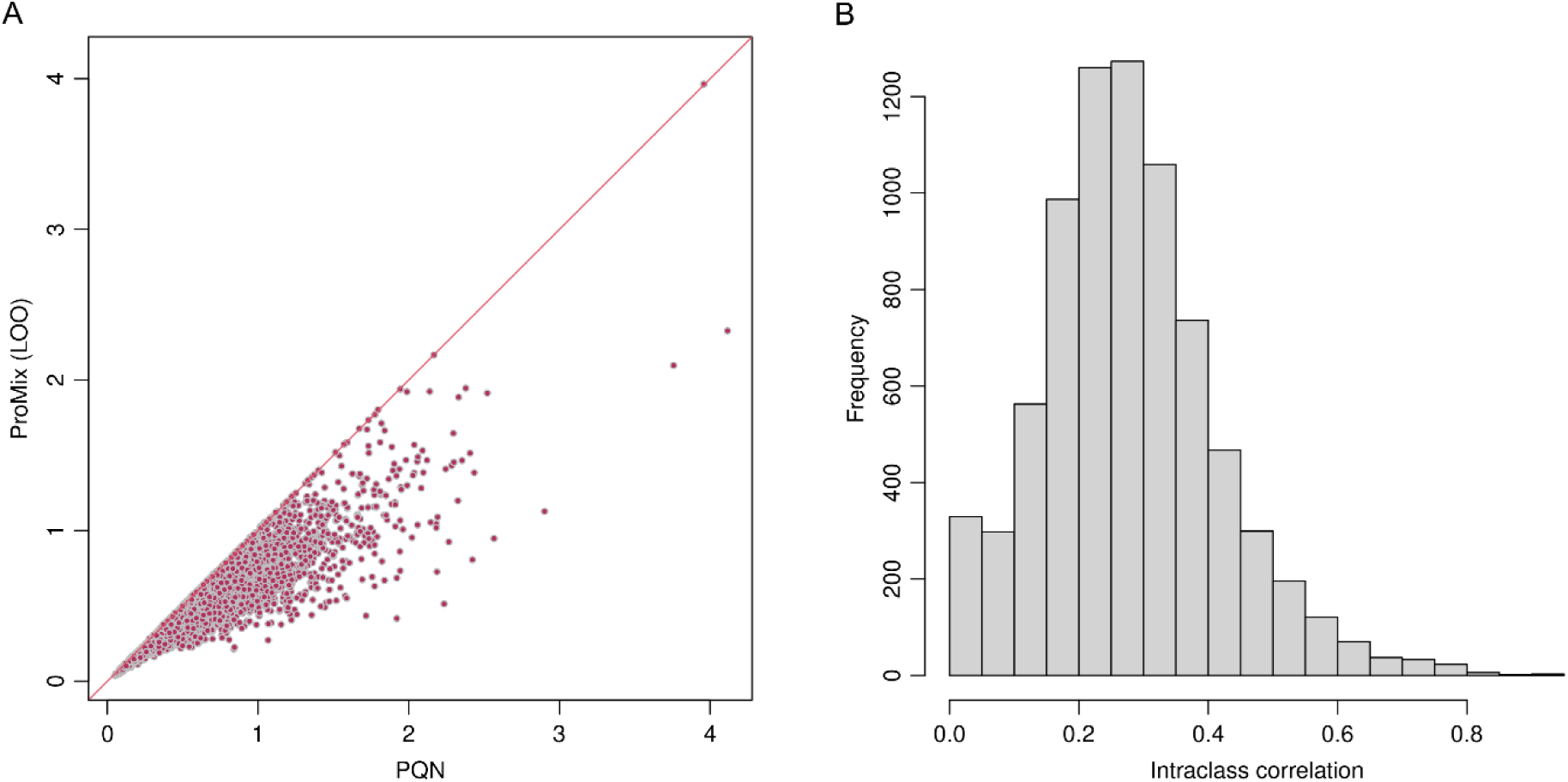
Normalization of GTEx lung data. (A) Mean squared error (repRMSE) between technical replicates using PQN (x-axis) and ProMix with leave-one-out (y-axis). (B) Histogram of the estimated intra-class correlation.

The right panel of Fig 6 displays the estimated intraclass correlation resulting from the run effects, defined as 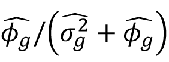. Across the 7762 proteins, the median intraclass correlation is 0.27 (the 25th- and 75th-percentiles are 0.19 and 0.35, respectively), suggesting a moderate correlation induced by the variability of the tissue-specific reference. Encouragingly, in a downstream mapping of protein quantitative trait loci (pQTL), 251 pQTLs were identified at FDR < 0.1 using ProMix normalization, compared to 216 using PQN, suggesting that ProMix more accurately captures the biological variability among specimens.

## Discussions

Recent and ongoing developments in MS-based proteomics approaches, such as TMT and iTRAQ, have rapidly advanced the scale and depth with which the identity, quantity, modification, localization, and interaction of proteins can be interrogated and compared. This study introduces and evaluates two experimental design considerations for enhancing the precision and consistency of protein quantification: the inclusion of technical replicates and the inclusion of a second reference sample in each run. Improving the consistency of protein quantification across MS runs, while reducing sampling variability, is particularly important in studies where the true variation in protein abundance is small compared to technical measurement error. A case in point is proteogenomic studies aimed at identifying pQTLs. With a sample size of over 50,000, a plasma-based study revealed that more than 80% of the 2922 proteins had at least one pQTL^13^; however, achieving such a sample size is unrealistic in most studies focusing on specific tissues, cell types, or diseases.

Here, we focus on reducing the sources of variation between MS runs, which not only represent a major source of variability but also create an undesirable correlation structure between samples within the same run (Fig 6B). Using an ongoing proteogenomic study of lung tissue as an example, we investigated the efficacy of two augmented experimental strategies: the inclusion of technical replicates and the inclusion of a second reference sample. Analyses of real and simulated data demonstrated that both strategies enable the estimation of run-level effects, which can be adjusted during the normalization step. Our simulation results indicate that technical replicates alone are effective for accounting for variability between runs as well as for reducing measurement variability for each specimen. However, when replicates are not feasible, for example due to limitation in the quantity of specimens, a second reference sample can be used to account run-specific effects. Compared to full replicates, which double the number of MS runs, adding a second reference is also more economical with respect to MS time and cost. Currently, up to 18 samples can be multiplexed in a single run; thus, the cost of including a second reference is less than 6% of the cost of a full run. We note that these two augmented experimental features are beneficial for purposes unrelated to normalization. In the GTEx lung study, the primary motivation for including technical replicates was to increase proteome coverage and reduce the proportion of missing data, while the primary motivation for including a second reference was to facilitate comparisons with data generated from a previous TMT-based proteomic study^8^.

We propose a linear mixed model, implemented in ProMix, for the normalization of multiplex proteomics experiments. ProMix aims to estimate and adjust for variability across runs by leveraging technical replicates and/or additional reference samples. We note that ProMix bears similarity to MSstats^14^ in that both adopt a linear mixed model. However, MSstats has been developed for testing differential protein abundance across conditions under a split-plot design. Subjects within each condition are typically treated as exchangeable units; hence the individual subject effect is modeled as a random effect and not explicitly estimated. In contrast, the goal of a proteogenomic study is to test the association between protein abundance and thousands of genetic variants; furthermore, subject-level covariates, such as age and sex, need to be adjusted in the analysis. Therefore, MSstats cannot be directly applied.

Analyses of simulated and real data demonstrate that ProMix can substantially improve the normalized abundance compared to existing methods, such as PQN. By implementing a coordinate-wise descent algorithm, ProMix is computationally efficient. We also show that repMSE computed from ProMix-LOO provides an informative diagnostic for the necessity and usefulness of ProMix. In our analysis of the GTEx lung study, we observed that PQN produced a lower repMSE than ProMix-LOO for approximately 6% of proteins. A lower repMSE produced by PQN may indicate either low run-level variability compared to other sources of stochasticity or that the run-specific effects are poorly estimated. For these proteins, one can simply choose to use PQN instead. Anecdotally, we found that some of these poorly performing proteins feature missing data from entire runs, while others have their mixed-effects model estimation strongly influenced by a few outlier observations or rendered numerically unstable. For these cases, a future direction is to model the underlying distribution that represents the run-level effects. This distribution could be specified as a Bayesian prior or estimated from data using an empirical Bayes approach, following a strategy adopted for modeling gene-specific dispersion factors in RNA-seq analysis^15^. Finally, the linear mixed model introduced in ProMix offers a flexible framework for incorporating emerging innovations in multiplex quantitative proteomic technology.

---

**Figure S1.**
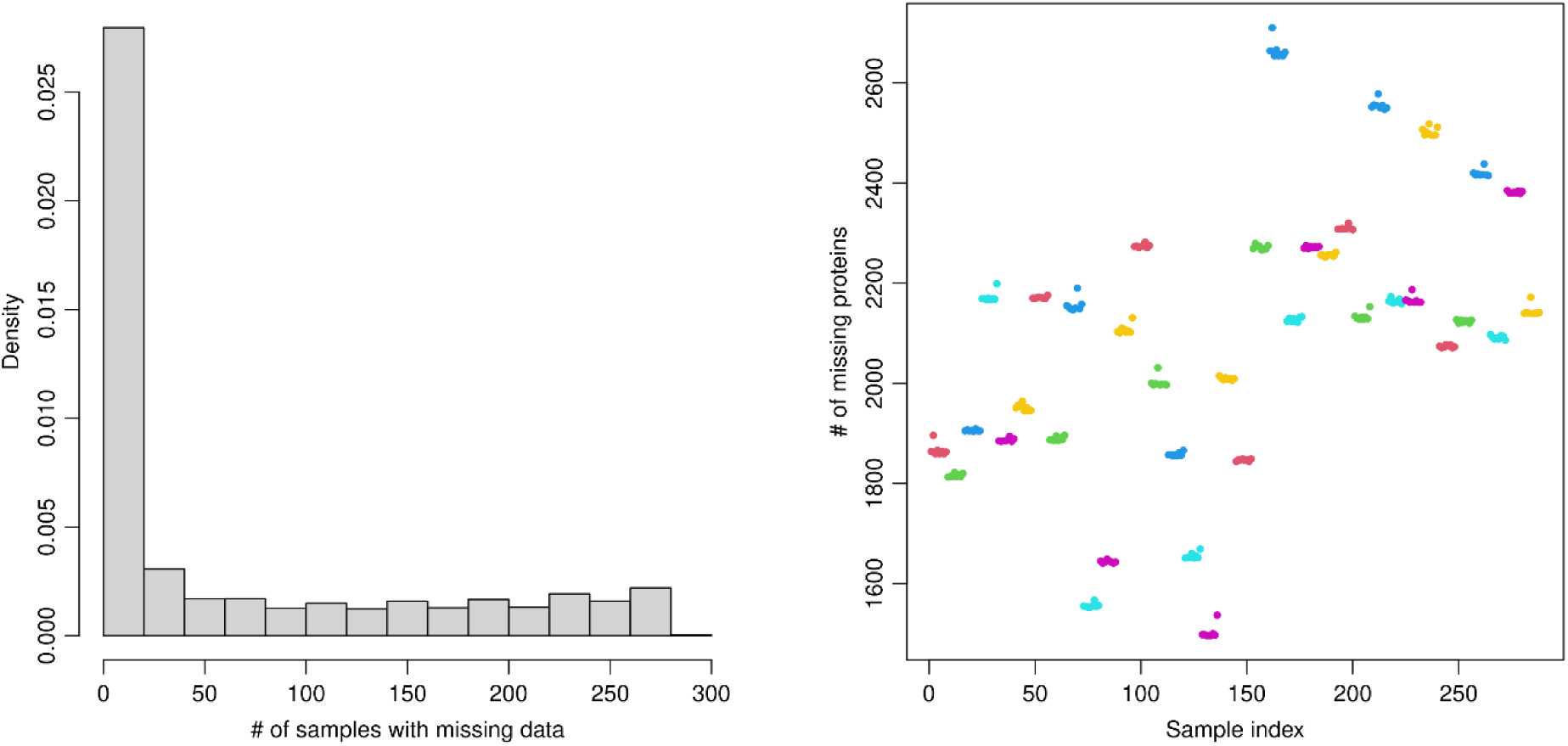
Missing pattern of lung data. (A) Protein-wise missing pattern. (B) Sample-wise missing pattern; samples are grouped in order of mass spectrometry runs. Each group of consecutive points with the same color delineates a mass spectrometry run. Note that the missing pattern is highly correlated between samples within a run.

**Figure S2.**
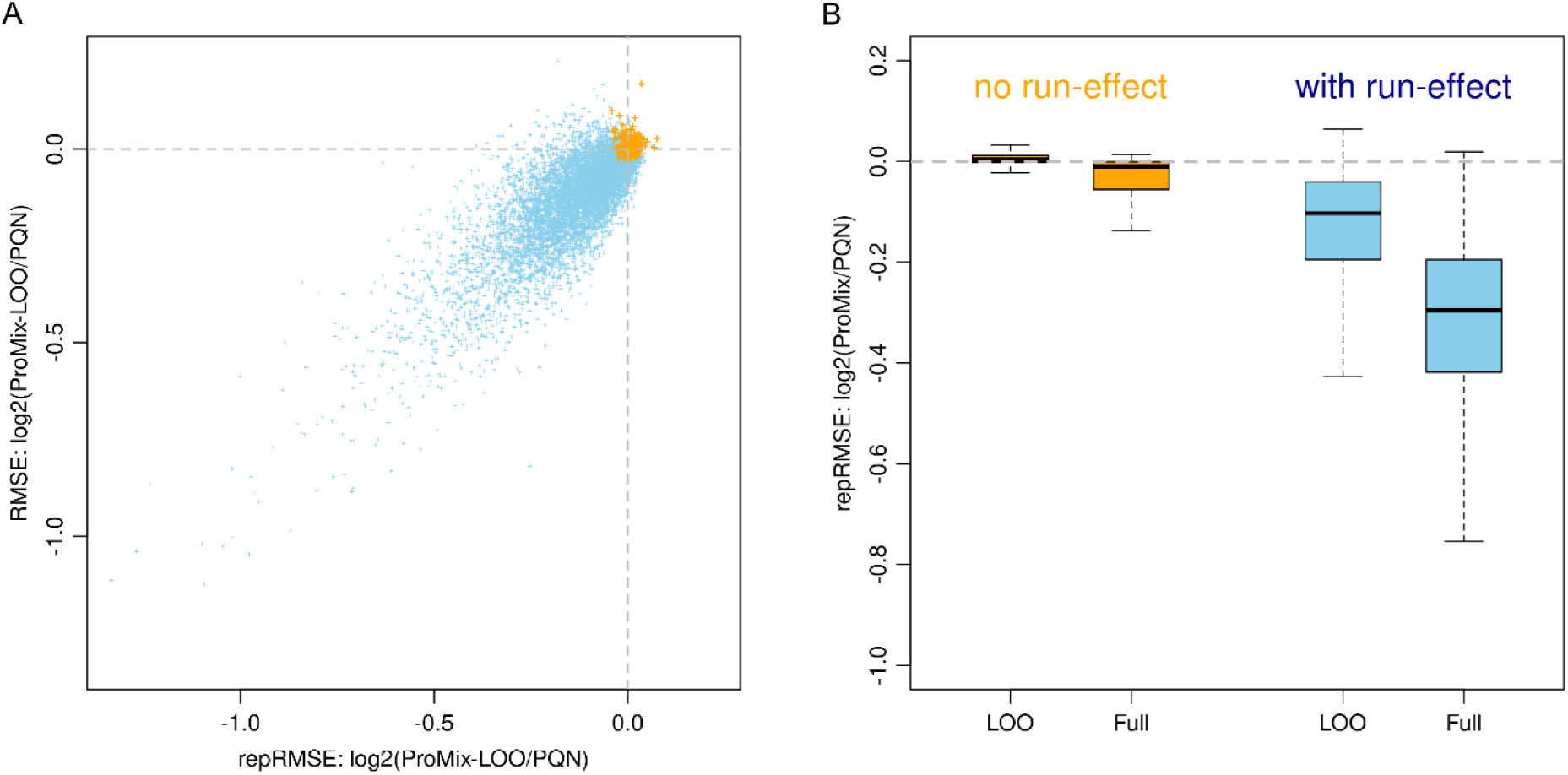
Using repRMSE to inform MSE. (A) Relationship between the reduction in RMSE within replications (repRMSE) and the reduction in RMSE to the true protein abundance. X-axis: 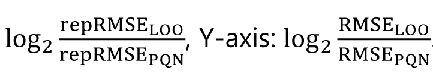 Each point represents a simulated protein. Blue points represent proteins with run effects (𝜙_𝑔_ > 0); orange points represent proteins without run effects (𝜙_𝑔_ = 0), for which PQN suffices. The general positive correlation pattern suggests that the reduction in repRMSE can serve as an informative surrogate for the reduction in RMSE between estimated and true protein abundance. (B) Boxplots of repRMSE computed by ProMix using leave-one-out (LOO) or full data mode. The Y-axis represents log_2_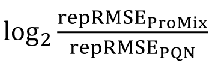 proteins without run effects (orange boxes), ProMix-LOO achieves comparable repRMSE to using PQN; in contrast, ProMix using full data obtains, on average, lower repRMSE due to overfitting.

## References

1. Ross, P. L. et al. Multiplexed protein quantitation in Saccharomyces cerevisiae using amine-reactive isobaric tagging reagents. Mol Cell Proteomics 3, 1154–1169 (2004).

2. Dayon, L. et al. Relative quantification of proteins in human cerebrospinal fluids by MS/MS using 6-plex isobaric tags. Anal Chem 80, 2921–2931 (2008).

3. Petralia, F. et al. Pan-cancer proteogenomics characterization of tumor immunity. Cell. 187, 1255–1277.e27 (2024).

4. Wu, L. et al. Variation and genetic control of protein abundance in humans. Nature 499, 79–82 (2013).

5. Wang, J. et al. Pan-Cancer Proteomics Analysis to Identify Tumor-Enriched and Highly Expressed Cell Surface Antigens as Potential Targets for Cancer Therapeutics. Mol Cell Proteomics 22, 100626 (2023).

6. Dieterle, F., Ross, A., Schlotterbeck, G. & Senn, H. Probabilistic quotient normalization as robust method to account for dilution of complex biological mixtures. Application in 1H NMR metabonomics. *Anal Chem* **78**, 4281–4290 (2006).

7. Ardlie, K. G. et al. The Genotype-Tissue Expression (GTEx) pilot analysis: Multitissue gene regulation in humans. Science 348, 648–660 (2015).

8. Jiang, L. et al. A Quantitative Proteome Map of the Human Body. Cell 183, 269–283.e19 (2020).

9. Kong, A. T., Leprevost, F. V., Avtonomov, D. M., Mellacheruvu, D. & Nesvizhskii, A. I. MSFragger: Ultrafast and comprehensive peptide identification in mass spectrometry-based proteomics. Nat Methods 14, 513–520 (2017).

10. da Veiga Leprevost, F., et al. Philosopher: a versatile toolkit for shotgun proteomics data analysis. Nat Methods 17, 869–870 (2020).

11. Li, G. X. et al. Comprehensive proteogenomic characterization of rare kidney tumors. Cell Rep Med 5, 101547 (2024).

12. Bates, D., Mächler, M., Bolker, B. M. & Walker, S. C. Fitting linear mixed-effects models using lme4. J Stat Softw 67, 1–48 (2015).

13. Sun, B. B. et al. Plasma proteomic associations with genetics and health in the UK Biobank. Nature 622, 329–338 (2023).

14. Kohler, D. et al. MSstats Version 4.0: Statistical Analyses of Quantitative Mass Spectrometry-Based Proteomic Experiments with Chromatography-Based Quantification at Scale. J Proteome Res 22, 1466–1482 (2023).

15. McCarthy, D. J., Chen, Y. & Smyth, G. K. Differential expression analysis of multifactor RNA-Seq experiments with respect to biological variation. Nucleic Acids Res 40, 4288– 4297 (2012).

